# 3D black blood MRI atlases of congenital aortic arch anomalies and the normal fetal heart: application to automated multi-label segmentation

**DOI:** 10.1101/2022.01.16.476503

**Authors:** Alena U. Uus, Milou P.M. van Poppel, Johannes K. Steinweg, Irina Grigorescu, Alexia Egloff Collado, Paula Ramirez Gilliland, Thomas A. Roberts, Mary A. Rutherford, Joseph V. Hajnal, David F.A. Lloyd, Kuberan Pushparajah, Maria Deprez

## Abstract

**Background:** 3D image-domain reconstruction of black blood contrast T2w SSTSE fetal MRI datasets using slice-to-volume registration methods showed to provide high-resolution 3D images of the heart with superior visualisation of fetal aortic arch anomalies [1]. However, there is a lack of formalisation of the MRI appearance of fetal cardiovascular anatomy and standardisation of vessel segmentation protocols.

**Methods:** In this work, we present the first set of 3D fetal MRI atlases defining normal and abnormal fetal aortic arch anatomy created from 3D reconstructed images from 87 subjects scanned between 29-34 weeks of gestation with postnatally confirmed outcomes. We also implement and evaluate atlas-guided registration and deep learning (UNETR) methods for automated 3D multi-label fetal heart vessel segmentation.

**Results:** We created four atlases representing the average anatomy of the normal fetal heart, coarctation of the aorta, right aortic arch and suspected double aortic arch. Inspection of atlases confirmed the expected pronounced differences in the anatomy of the aortic arch. The results of the multi-label heart vessel UNETR segmentation showed 100% per-vessel detection rate for both normal and abnormal aortic arch anatomy.

**Conclusions:** This work introduces the first set of 3D black blood T2w MRI atlases of the normal and abnormal fetal cardiovascular anatomy along with detailed segmentation of the major cardiovascular structures. We also demonstrated the feasibility of using deep learning for multi-label vessel segmentation.

## Background

The black blood contrast in T2w single shot turbo spin echo (SSTSE) sequence widely used in fetal MRI provides superior visualisation of fetal cardiovascular anomalies [2]. However, 2D-only slice-wise analysis is generally characterised by low reliability for the small, complex 3D structures of the fetal cardiovascular system due to uncontrolled fetal motion during acquisition. Recently, application of slice-to-volume registration (SVR) motion correction tools [3, 4] for reconstruction of high-resolution 3D T2w images of the fetal heart was shown to provide adjunct diagnostic information and allow visualisation of the fetal extracardiac vasculature using 3D models [1].

At our institution, the main clinical referral indications for 3D T2w fetal cardiac MRI (CMR) [1] include suspected coarctation of the aorta (CoA), right aortic arch (RAA) with aberrant left subclavian artery (ALSA) and suspected double aortic arch (DAA). However, despite the reported improved diagnostic performance of 3D fetal CMR [1, 5], there is a lack of formalization of the MRI appearance of fetal cardiovascular anatomy. Furthermore, the currently employed 3D semi-automatic segmentation of the heart vasculature from T2w SVR images [1] requires time-consuming manual refinement, especially for multiple labels and is vulnerable to inter-observer bias.

### Contributions

In this work, we present the first 3D fetal MRI atlases defining normal and abnormal (suspected CoA, RAA+ALSA, DAA) fetal cardiovascular anatomy generated from 87 fetal CMR datasets along with the formalised multi-label parcellations of the major cardiovascular structures. This could potentially contribute to standardisation of the 3D SVR-based diagnostic protocol and quantitative analysis. We also perform evaluation of the feasibility of atlas-guided registration and deep learning methods for automated 3D multi-label vessel segmentation.

## Methods

### Datasets and preprocessing

The fetal MRI data include 87 datasets acquired under the iFIND [6] project at St. Thomas’s Hospital, London [REC: 14/LO/1806] and as a part of the fetal CMR service at Evelina London Children’s Hospital, with data use for this project subject to informed consent of the participants [REC: 07/H0707/105].

The cohorts (Fig. 1A) include 29 fetuses without reported cardiac anomalies (“normal”), 58 cases with postnatally confirmed abnormality (20 CoA, 21 RAA and 17 DAA). The inclusion criteria were high SNR, good visibility of extra-cardiac vasculature, singleton pregnancy, 29-34 weeks gestational age (GA) and absence of other cardiovascular anomalies or significant anatomical deviations. The acquisitions were performed on a 1.5T Philips Ingenia MRI system using torso receiver array and T2w SSTSE: TR=15000ms, TE=80ms, voxel size 1.25×1.25×2.5mm, slice thickness 2.5mm and spacing 1.25mm with 9-11 stacks. The 3D reconstructions of the fetal thorax (0.75mm isotropic resolution, standard radiological space) were generated using our new automated deformable SVR (DSVR) [7] reconstruction pipeline [4].

**Figure 1.**
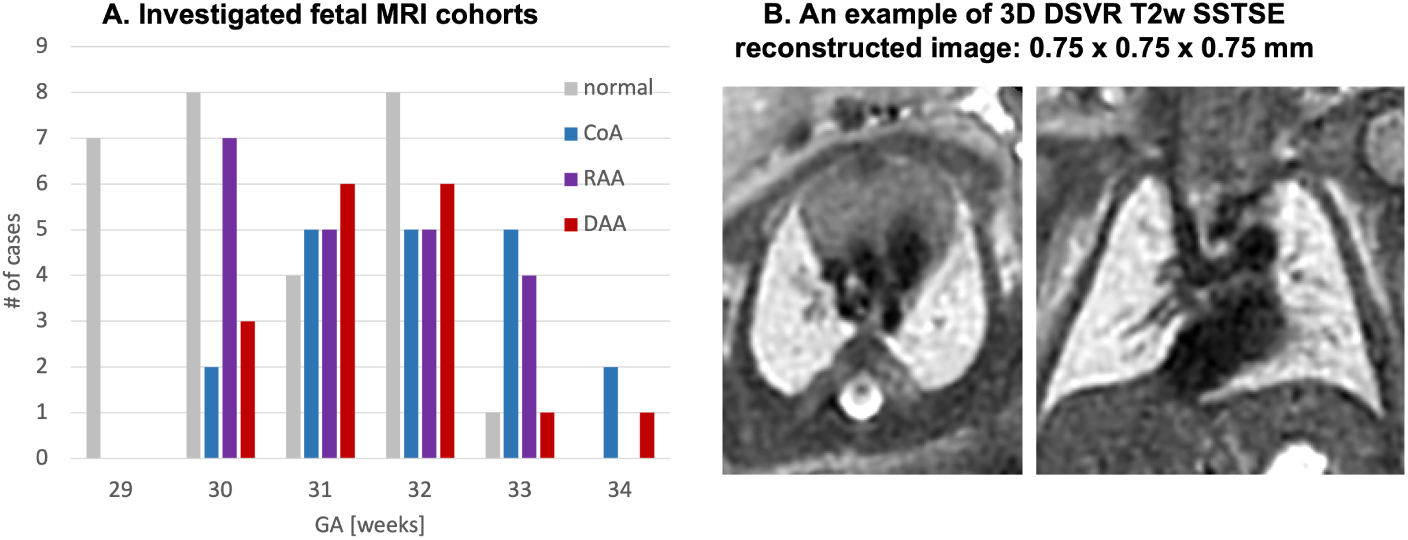
Investigated fetal MRI cohorts (A) and an example of 3D DSVR reconstructed thorax for one of the black blood T2w SSTSE datasets (B).

### Generation of 3D heart atlases with vessel segmentation

The 3D atlases for each of the groups (normal, CoA, RAA+ALSA, DAA) were created using the classical MIRTK atlas generation tool [8, 9] (Fig. 2). As preprocessing, the 3D reconstructed images from all cohort were affinely registered to the same standard space. The output atlases have 0.6mm isotropic resolution.

**Figure 2.**
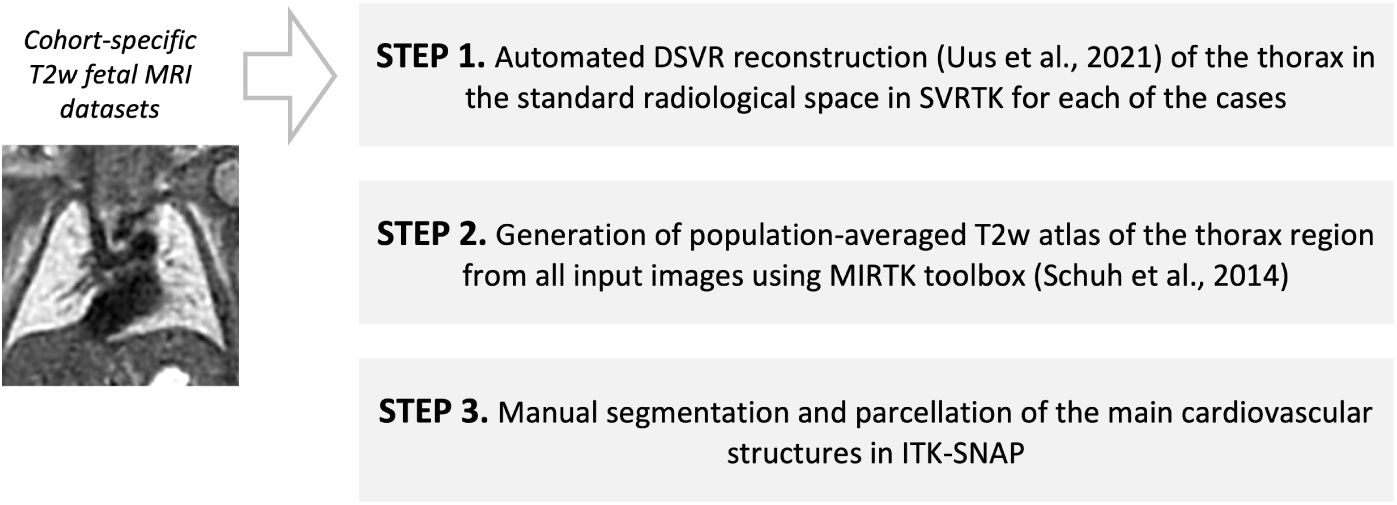
Main steps of the fetal heart atlas generation pipeline.

For each of the atlases, a clinician (MvP) trained in fetal CMR manually segmented parcellations of 20 main cardiovascular structures in ITK-SNAP [10]. The chosen structures follow the standard (fetal) CMR reporting protocol established in [1]and currently used in clinical practice. The atlases are publicly available online at SVRTK data repository [11].

### Automated segmentation of the fetal heart vessels

Fig. 3 shows the proposed pipeline for automated multi-label 3D segmentation of the cardiovascular structures in 3D DSVR MRI images. It was implemented in Pytorch MONAI framework [12] based on of the 3D UNETR transformer-based convolutional neural network (CNN) [13] that reportedly produce superior performance for multi-label segmentation.

**Figure 3.**
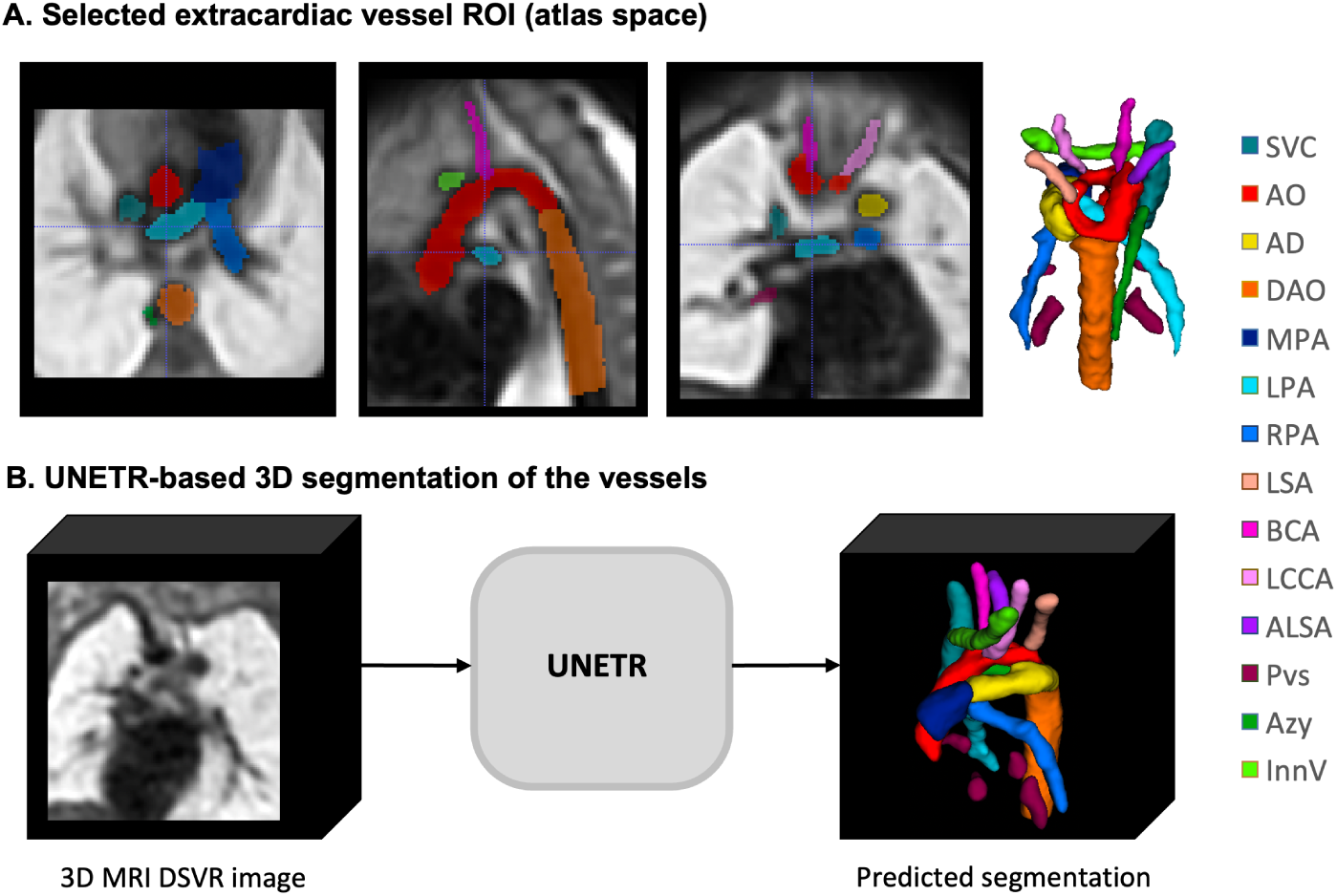
Definition of the vessels in the extracardiac ROI (A) and UNETR-based segmentation.

We used images from CoA, RAA and DAA cohorts including: 42 cases for training (14 from each cohort), 3 for validation and 6 for testing. The selection criteria were the isolated specific anomaly, absence of anatomical variations and the clear visibility and continuity of all vascular structures. The preprocessing included cropping to the extracardiac vessel ROI (e.g., Fig. 3A), affine registration to the atlas space and resampled with padding to 128×128×128 grid. The labels (14 different vessel ROIs) were generated using MIRTK registration-guided label propagation cohort-specific atlas label propagation (LP). This was followed by manual refinement of individual structures in all segmentations using ITK-SNAP, primarily for the head and neck vessels, azygous vein and aortic arch due to the limited visibility and partial volume effect. The RAA and DAA cases reportedly required more refinement than CoA due to the higher variations in the anatomy. All labels were visually assessed and confirmed to be qualitatively acceptable. The training was performed for 50000 iterations with the standard MONAI-based augmentation (random bias field, contrast adjustment, Gaussian noise and affine rotations ±45°). The performance was tested on 2 CoA, 2 RAA and 2 DAA cases vs. manually refined propagated labels in terms of the vessel detection status (visual assessment: correct=100%, partial=50%, failed=0%), sensitivity, specificity and Dice.

## Results

### 3D heart atlases

Fig. 4-6 show the generated 3D T2w atlases along with the corresponding multilabel parcellation maps. The anatomical accuracy of the atlases and the segmented structures were confirmed by a clinician with more than 4 years of fetal CMR experience. Visual assessment of the atlases (e.g., Fig. 4A) demonstrates high contrast of the vessels as well as the structural continuity of the blood pool. As expected, the image quality is higher than in the individual cases.

**Figure 4.**
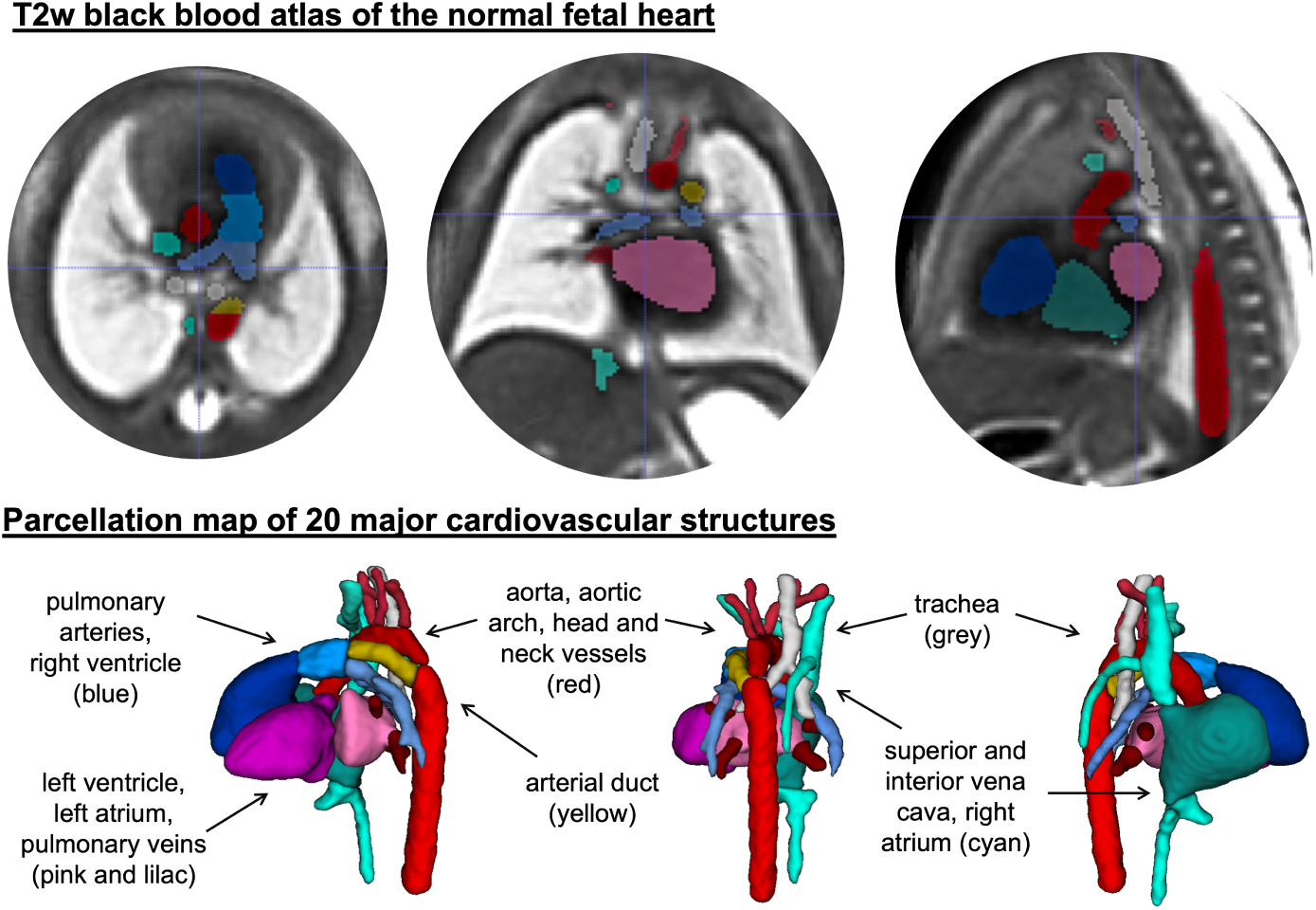
Generated atlas of the normal heart anatomy with the corresponding multi-label parcellation map.

The parcellation maps (e.g., Fig. 4B) include the four heart chambers and 14 major vascular structures analysed during the standard 3D fetal MRI-based diagnostic assessment [1] including: aorta and head and neck vessels (red), arterial duct (yellow), vena cava and innominate vein (cyan), pulmonary arteries (light blue) and pulmonary veins (lilac). The trachea is shown in grey.

The 3D model of the CoA atlas shown in Fig. 5A has a pronounced narrowing of the aortic arch in comparison to the normal anatomy. This is confirmed by the significantly smaller aortic arc diameter along the cenreline presented in Fig. 5B. The extraction of vessel diameter was performed in VMTK toolbox [14]. Importantly, the CoA atlas represents an averaged state of all CoA types [5] as the input cases had varying position and degree of arch narrowing.

**Figure 5.**
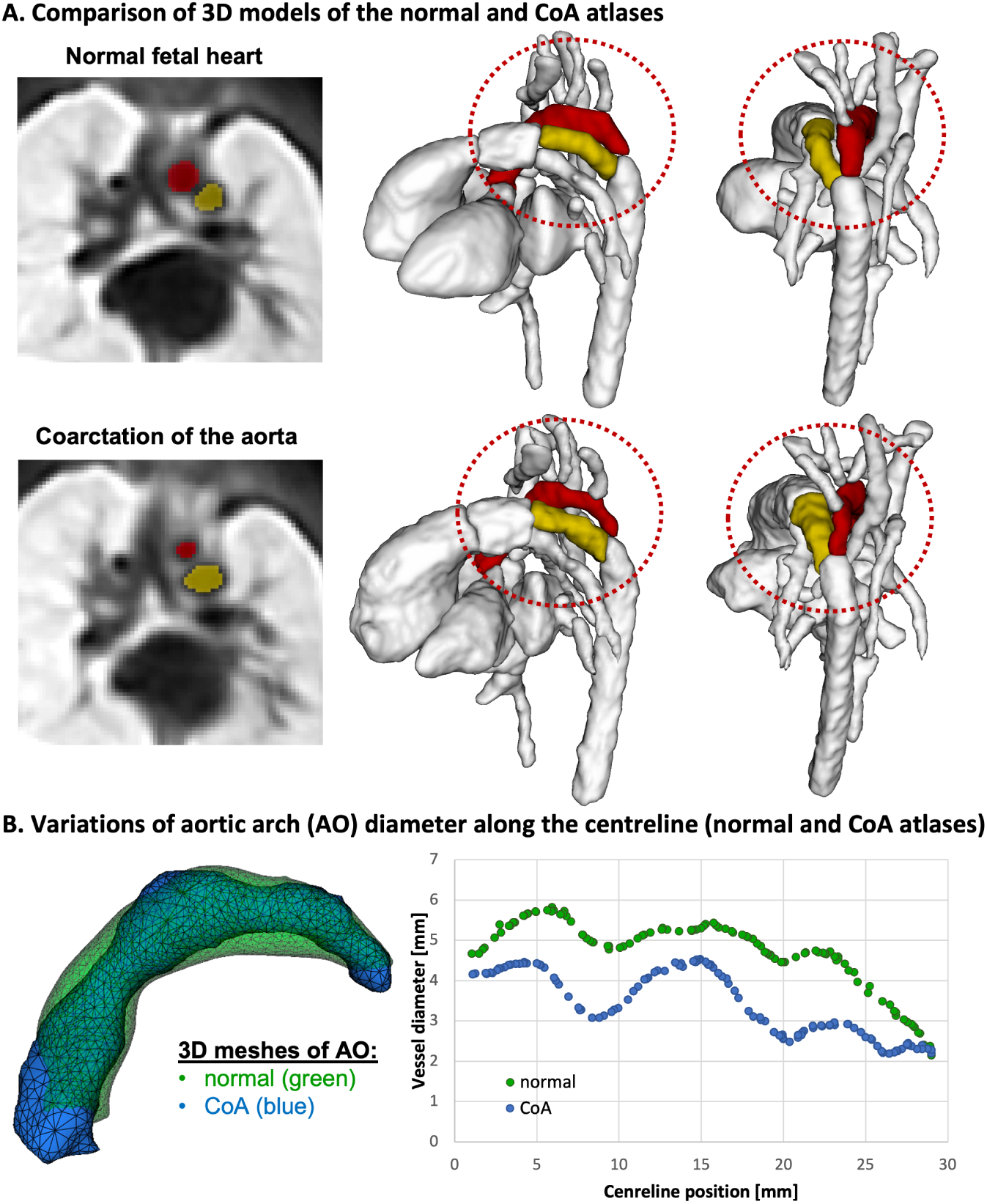
Comparison between the normal and abnormal CoA anatomy in the generated 3D atlases. A: The 3D models show narrowing of the aortic arch (red) in the CoA atlas. The arterial duct is visualised in yellow and the rest the anatomy is set to white. B: Variations in the normal (green) and CoA (blue) AO diameter along the cenreline.

For the RAA and DAA atlases, Fig. 6 clearly shows the expected [15] differences in the relative position of the trachea and the aortic arch(es) and aortic branching vs. the normal anatomy. Both RAA and DAA atlases also have the additional aberrant left subclavian artery (ALSA) highlighted in green.

**Figure 6.**
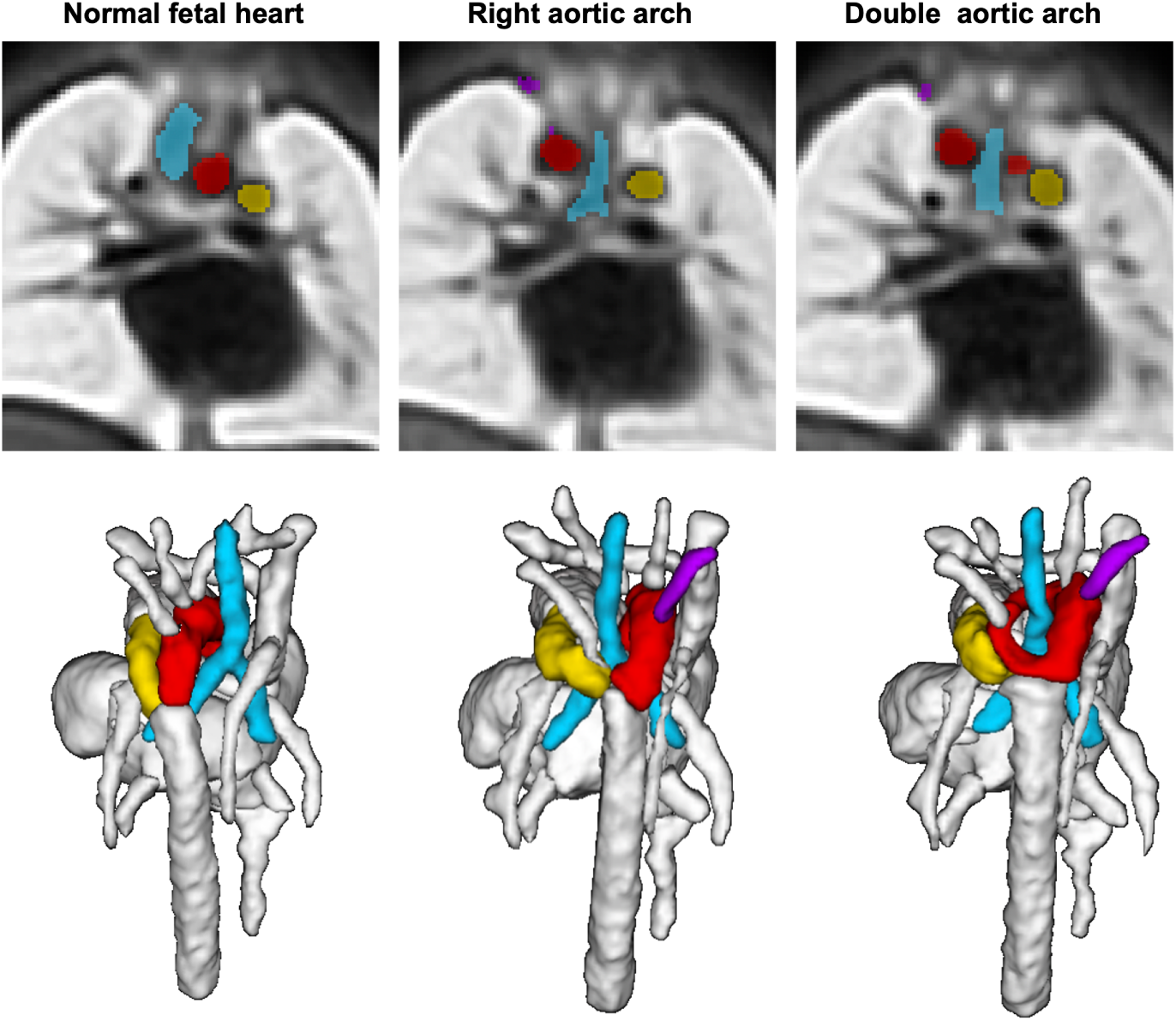
Comparison between the normal and abnormal RAA and DAA anatomy in the generated 3D atlases: differences in the relative position of the trachea (light blue) and aortic arch (red). The arterial duct is visualised in yellow and the ALSA is visualised in lilac.

### Automated heart vessel segmentation

The results of testing of the trained multi-label 3D UNETR segmentation network on 6 CoA, RAA and DAA cases are summarised in Fig. 7. The UNETR CNN correctly detected all vessels in all test subjects (100%) with different anomalies which is confirmed by high specificity and relatively high Dice values. Notably, even after manual refinement, the reference labels used in training and testing are prone to inconsistencies due to the the partial volume effect and varying per-vessel intensity contrast levels and cannot be considered as an absolute ground truth. This, along with the small vessel size and no clear signal definition along the entire length of the vessel contributed to the lower Dice and sensitivity values for certain vessels (e.g., BCA or LLCA). Nonetheless, these results confirm the feasibility of using CNN for multi-label segmentation for small vascular structures in a mixture of abnormal datasets with different anatomy.

**Figure 7.**
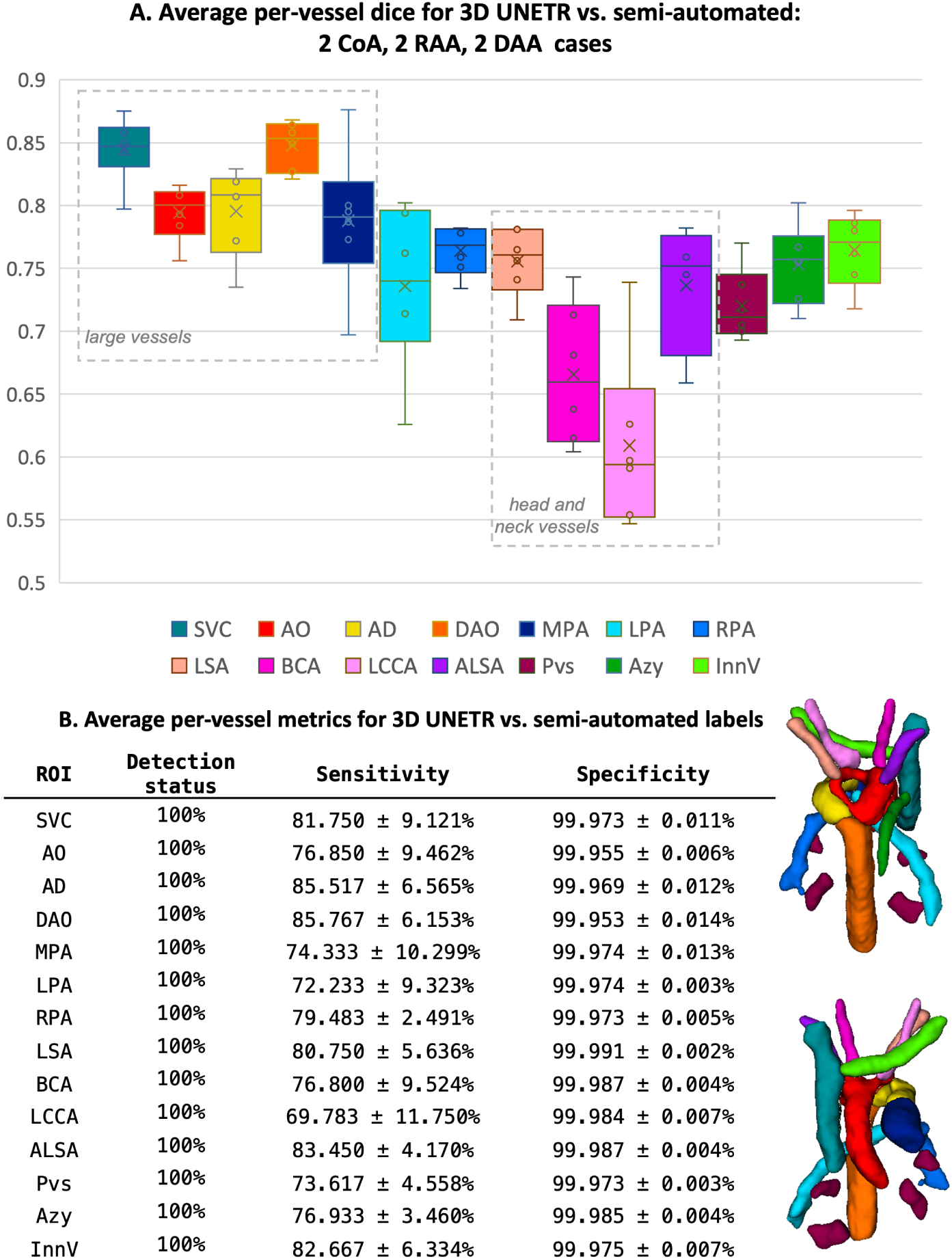
Evaluation of 3D UNETR segmentation for 14 extracardiac vessel ROIs including average Dice (A), sensitivity, specificity and detection detection status (B).

The examples of visual comparison of the ground truth manually refined labels propagated from the atlases vs. CNN outputs for different anomaly groups are shown in Fig. 8. While label propagation provided sufficiently good localisation of the vessels, there are present minor inconsistencies and patchy appearance of the labels at the vessel boundaries. This is primarily related to the expected limitations of intensity-based registration methods. These irregularities were partially resolved in the CNN output, which produced significantly more realistic and smoother boundaries and potentially corrected minor errors.

**Figure 8.**
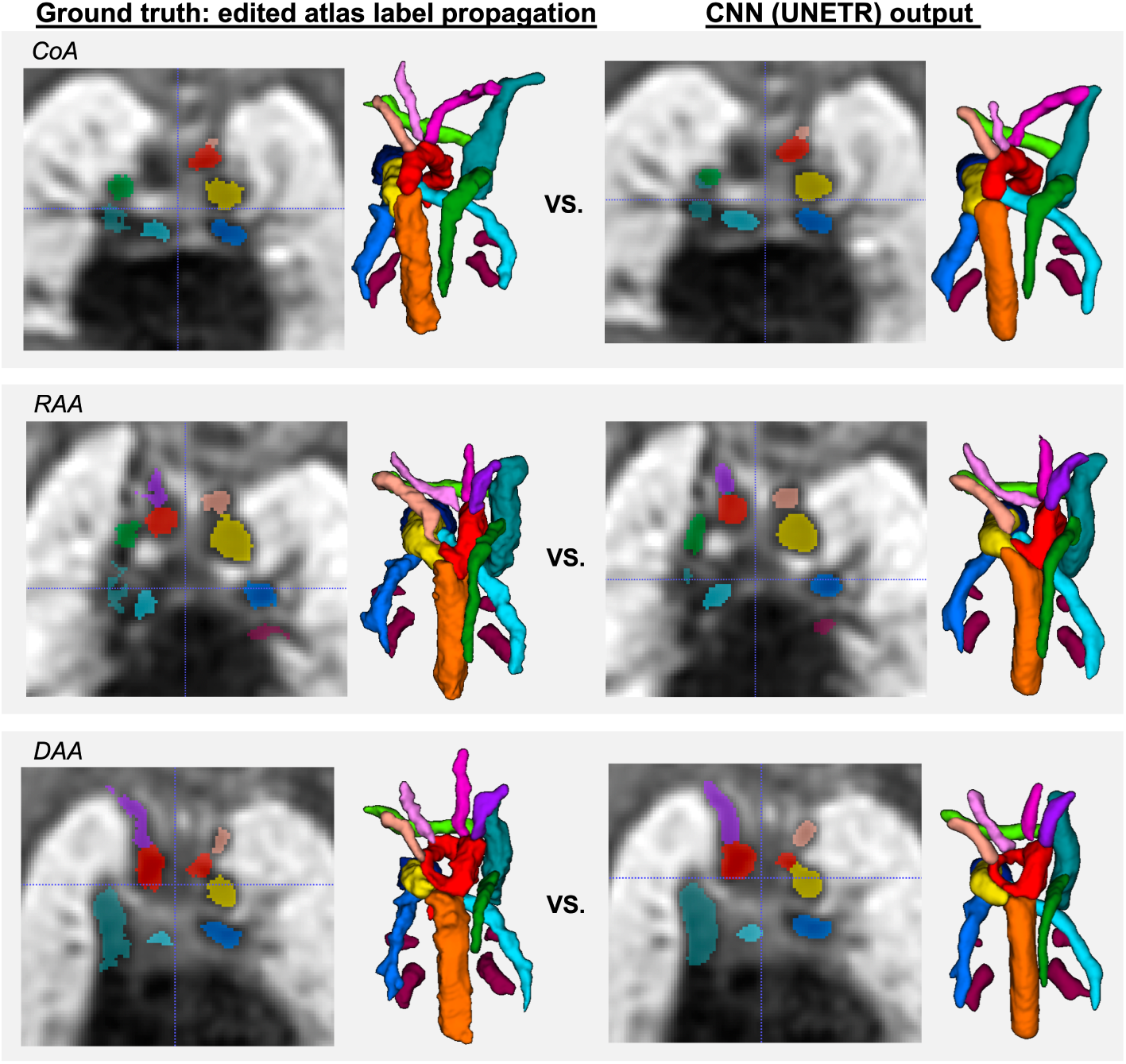
Examples of comparison between the manually refined propagated labels (ground truth) and CNN outputs for three different aortic arch anomaly groups.

This also confirms the feasibility of using labels propagated from average atlases for CNN training, which significantly reduces required dataset preparation time vs. manual-only segmentations. However, even without extreme anatomical deviations, label propagation outputs tend to require a certain amount of manual editing. E.g., Fig. 9 shows an example of failed label propagation segmentation of the azygous vein for one of the test datasets. It required manual correction while the CNN produced correct segmentation. In total, for all datasets used in training and testing of the networks, minor or significant manual editing was required in 30.2% and 10.1% of all individual structures, correspondingly.

**Figure 9.**
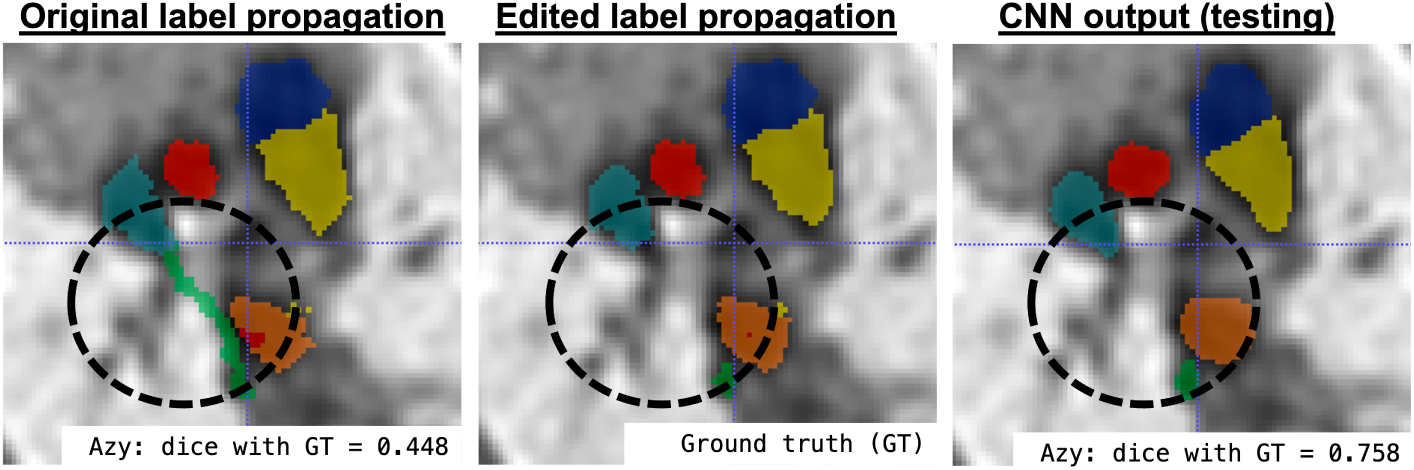
An example of label propagation error in one of the testing datasets that required manual refinement: incorrect azygous vein segmentation.

## Discussion

The proposed population-averaged black blood atlases of the normal and abnormal fetal heart anatomy define the expected appearance of cardiovascular structures in fetal MRI. The correctness of the vascular anatomy in all atlases was confirmed by experienced fetal CMR clinicians. The atlases are valuable as a general educational material and can also be used for parental counseling and diagnosis.

Another application of the atlases is label propagation for multi-vessel segmentation. However, while it produces relatively consistent results for the normal or close-to-normal anatomy in CoA cases, DAA and RAA cases are the subject to higher anatomical variations, which leads to lower segmentation quality. Varying image quality is another aspect that impedes expected segmentation quality in registration-based methods. The modern advanced CNN-based segmentation methods provide an alternative operational solution but they also require large and high quality training datasets.

In this work, we utilised manually refined atlas-based segmentations for CNN training, which significantly reduced time for preparation of the datasets and provided additional consistency of the labels. The results of testing confirmed that a single conventional CNN can be successfully used for multi-label segmentation of abnormal datasets with different aortic arch anatomy.

In the context of 3D MRI-based congenital CHD diagnosis, the main aims of vessel segmentation is visualisation of the cardiac vasculature in 3D for confirmation of the presence and assessment of the relative position and dimensions of all relevant structures. This is especially important for differentiation between RAA, DAA and the normal positioning of the large vessels. Additionally, diagnosis of suspected CoA would require quantitative measurements of the diameter variations and angles of the aortic arch that showed to correlate with the neonatal outcomes [5]. On the other hand, assessment of the smaller vessels would primarily need only the detection status due to the lower reliability, visibility and image quality. In general, even with the lower dice for small vessels, the results of this work confirm the feasibility of the proposed approach for fetal heart vessel segmentation.

One of the main requirements to any segmentation-based image analysis on is the consistency segmentation protocols. Potentially, this can be achieved by deep learning since a one common segmentation network could potentially reduce inter-observer bias for both visualisation and quantitative measurements including general biometry and shape analysis.

However, the reliability of segmentation results for routine clinical cases would depend on the degree of anatomical variability and presence of other anomalies in training datasets, which should also include different MRI acquisition protocols. Therefore, a universal solution will require preparation of a significantly larger training dataset as well as implementation of quantification of uncertainty of segmentation of individual structures.

## Conclusions

This work introduced the first set of 3D black blood T2w fetal MRI heart atlases of the normal and abnormal aortic arch anatomy (CoA, RAA, DAA) along with detailed parcellation maps of the major cardiovascular structures. This is a first step towards standardisation of the analysis and diagnosis protocol for fetal cardiovascular anomalies using 3D motion-corrected MRI [1].

In addition, we demonstrated the feasibility of using deep learning for automated multi-label vessel segmentation for different types of aortic arch arch anomalies.

Our future work will focus on optimisation of the CNN-based segmentation pipeline for a wider range of fetal cardiovascular abnormalities and anatomical variations as well as different acquisition protocols and automated biometry.

## Acknowledgements

We thank everyone who was involved in acquisition and examination of the datasets and all participating mothers. The views expressed are those of the authors and not necessarily those of the NHS, the NIHR or the Department of Health

## Funding

This work was supported by the Rosetrees Trust [A2725], the Wellcome/EPSRC Centre for Medical Engineering at King’s College London [WT 203148/Z/16/Z], the Wellcome Trust and EPSRC IEH award [102431] for the iFIND project, the NIHR Clinical Research Facility (CRF) at Guy’s and St Thomas’ and by the National Institute for Health Research Biomedical Research Centre based at Guy’s and St Thomas’ NHS Foundation Trust and King’s College London.

## Abbreviations

DAA: double aortic arch
AD: arterial duct
AO: aorta
ALSA: aberrant left subclavian artery
Azy: azygous vein
BCA: brachiocephalic artery
CoA: coarctation of the aorta
CHD: congenital heart disease
CNN: convolutional neural networks
CMR: cardiac magnetic resonance imaging
DAO: descending aorta
DSVR: slice-to-volume registration
InnV: Innominate vein
IVC: inferior vena cava
MPA: main pulmonary artery
MRI: magnetic resonance imaging
LA: left atrium
LCCA: left common carotid artery
LPA: left pulmonary artery
LSA: left subclavian artery
LV: left ventricle
Pvs: pulmonary veins
RA: right artrium
RAA: right aortic arch
RPA: right pulmonary artery
RV: right ventricle
SSTSE: single shot turbo spin echo
SVC: superior vena cava
SVR: slice-to-volume registration

## Availability of data and materials

The generated 3D fetal heart MRI atlases with parcellation maps are publicly available at the SVRTK data repository [11].

## Ethics approval and consent to participate

All fetal MRI datasets used in this work were processed subject to informed consent of the participants [REC: 07/H0707/105; REC: 14/LO/1806].

## Competing interests

The authors declare that they have no competing interests.

## Consent for publication

All authors gave final approval for publication and agree to be held accountable for the work performed therein.

## Authors’ contributions

AU prepared the manuscript, processed datasets, implemented the code for segmentation, generated the atlases and conducted the experiments. MvP designed multi-label heart parcellation protocol, segmented the atlases and analysed the results. MvP and JK participated in data acquisition and processing of the datasets. JK participated in data acquisition and processing of the datasets. IG and PRG participated in implementation of the code for segmentation and analysis of the results. TR participated in data acquisition. JvH and MR are the leaders of the iFIND project. DL and KP formalised fetal CMR MRI acquisition protocol and SVR-based diagnosis protocol and and participated in analysis of the results. MD and KP conceptualized the study and the methods, obtained the funding and supervised all stages of the research and preparation of the manuscript.

## Notes

https://gin.g-node.org/SVRTK/fetal_mri_atlases

### Competing Interest Statement

The authors have declared no competing interest.

